# Identification of effector metabolites using exometabolite profiling of diverse microalgae

**DOI:** 10.1101/2021.07.02.450979

**Authors:** Vanessa Brisson, Xavier Mayali, Benjamin Bowen, Amber Golini, Michael Thelen, Rhona K. Stuart, Trent R. Northen

## Abstract

Dissolved exometabolites mediate algal interactions in aquatic ecosystems, but microalgal exometabolomes remain understudied. We conducted an untargeted metabolomic analysis of non-polar exometabolites exuded from four phylogenetically and ecologically diverse eukaryotic microalgal strains grown in the laboratory: freshwater *Chlamydomonas reinhardtii*, brackish *Desmodesmus* sp., marine *Phaeodactylum tricornutum*, and marine *Microchloropsis salina*, to identify released metabolites based on relative enrichment in the exometabolomes compared to cell pellet metabolomes. Exudates from the different taxa were distinct, but we did not observe clear phylogenetic patterns. We used feature based molecular networking to explore the identities of these metabolites, revealing several distinct di- and tripeptides secreted by each of the algae, lumichrome, a compound that is known to be involved in plant growth and bacterial quorum sensing, and novel prostaglandin-like compounds. We further investigated the impacts of exogenous additions of eight compounds selected based on exometabolome enrichment on algal growth. Of the these, five (lumichrome, 5’-S-methyl-5’-thioadenosine, 17-phenyl trinor prostaglandin A2, dodecanedioic acid, and aleuritic acid) impacted growth in at least one of the algal cultures. Two of these (dodecanedioic acid and aleuritic acid) produced contrasting results, increasing growth in some algae and decreasing growth in others. Together, our results reveal new groups of microalgal exometabolites, some of which could alter algal growth when provided exogenously, suggesting potential roles in allelopathy and algal interactions.

**IMPORTANCE:** Microalgae are responsible for nearly half of primary production on earth and play an important role in global biogeochemical cycling as well as in a range of industrial applications. Algal exometabolites are important mediators of algal-algal and algal-bacterial interactions that ultimately affect algal growth and physiology. In this study we characterize exometabolomes across marine and freshwater algae to gain insights into the diverse metabolites they release into their environments (“exudates”). We observe that while phylogeny can play a role in exometabolome content, environmental conditions or habitat origin (freshwater vs marine) are also important. We also find that several of these compounds can influence algal growth (as measured by chlorophyll production) when provided exogenously, highlighting the importance of characterization of these novel compounds and their role in microalgal ecophysiology.

## INTRODUCTION

Microalgae are important contributors to global carbon cycling, contributing almost 50% of photosynthetic carbon fixation (1). They are also important industrially for the production of biofuels and bioproducts (2, 3). The functionality and stability of algal systems, both natural and engineered, are influenced by interactions of algae with other microorganisms and with each other, and exometabolites released into the water are important mediators of these interactions (4).

The specific compositions of algal exometabolite pools are important for modulating algal growth, and the interactions of algae and their associated communities through nutrient exchange, signaling, growth promotion and inhibition, and defense. A range of different plant hormones (indoel-3-acetic acid, gibberellic acid, kinetin, 1-triacontanol, and abscisic acid) have all been shown to influence the growth of the model green alga *Chlamydomonas reinhardtii* (5). One strain of the coccolithophore *Emiliana huxleyi* has been shown to produce the plant hormone indole-3-acetic acid, while the growth of another strain was impacted by the same metabolite (6). Some algae have also been shown to inhibit the growth of competitor algae through the release of allelochemicals (7). Algal exudates have been hypothesized to play similar roles in the phycosphere, the microenvironment around algal cells, to those of root exudates in the rhizosphere (8). In plant systems, specific metabolites in root exudates have been shown to impact the growth of different microorganisms in the rhizosphere, favoring some microbial taxa over others (9). Similarly, combinations of metabolites known to be produced by algae have been shown to modulate microbial community composition depending on the combination of metabolites (10). In another study, rosmarinic acid and azelaic acid were shown to be produced by the diatom *Asterionellopsis glacialis* in response to bacterial presence and were also found to impact bacterial growth (11). Dimethylsulfoniopropionate produced by *E. huxleyi* feeds the bacterium *Phaeobacter inhibens*, but when the alga begins to produce p-coumarate, the bacterium responds by shifting its role from a mutualist, promoting algal growth, to a parasite, through the release of exometabolites (12).

Most studies of algal metabolomics have focused on the composition of intracellular metabolomes. For instance, of the organisms used in our study, the intracellular metabolomes of *Chlamydomonas reinhardtii* and *Phaeodactylum tricornutum* have been characterized for their response to different growth conditions (13, 14). In contrast, microalgal exometabolomes, particularly for marine microalgae, have not been well characterized, partly due to the technical challenges of low metabolite concentrations and high salt content in the samples (15). Some recent studies have begun to analyze algal exometabolomes: Shibil et al. investigated the exometabolome of the diatom *Asterionellopsis glacialis*, and its response to the addition of a microbial community (11), while Ferrer-Gonzáles et al. characterized the exometabolome of the diatom *Thalassiosira pseudonona* in comparison to co-cultures of the diatom with three different bacterial strains (15). Another study used an untargeted metabolomics analysis to compare the exometabolomes of cyanobacteria (three *Prochlorococcus* strains and two *Synechococcus* strain) to diatoms (two *Thalassosira* species and one *Phaeodactylum*) (16). Metabolite exudation and uptake have also been studied in the cyanobacterium *Synechococcus* sp. PCC 7002 (17). To better understand microalgal interactions, there is a need to characterize exometabolites from a broader range of microalgae.

In this study we characterized the non-polar exometabolomes of four phylogenetically and ecologically diverse eukaryotic microalgal strains: the freshwater green alga *Chlamydomonas reinhardtii*, the brackish green alga *Desmodesmus* sp. C046, the marine diatom *Phaeodactylum tricornutum*, and the marine eustigmatophycean alga *Microchloropsis salina* (also referred to as *Nannochloropsis salina*). We chose these strains to cover a phylogenetically diverse group of microalgae with different environmental origins. *C. reinhardtii* and *Desmodesmus* are the most closely related of these algae, both belonging to the plant kingdom in the phylum Chlorophyta and the class Chlorophyceae. *P. tricornutum* and *M. salina* are both members of the kingdom Chromista but belong to different phyla. Additionally, *C. reinhardtii* and *P. tricornutum* are both model organisms that have been well studied and whose genomes have been sequenced (18, 19). *Desmodesmus* is also of particular interest because it can grow in both freshwater and saltwater, allowing us to examine metabolomic shifts under these two conditions. In addition to profiling the exometabolomes, the goal of our analysis was to determine whether specific exometabolites detected from these strains impacted algal growth and might therefore be effector metabolites involved in interactions within or between algal strains.

## RESULTS

### Algal growth

Algal chlorophyll fluorescence, an indicator of algal growth, increased with incubation time for all algal strains through the eighth day after inoculation, when samples were collected for metabolomic analysis (Figure 1). Chlorophyll A fluorescence began to decrease by day 12 for *P. tricornutum*, and by day 18 for all algal cultures except for *C. reinhardtii*, indicating that all cultures were in mid to late exponential growth when samples were collected. Ash free dry weight measurements were made for the sampling times before and after day eight (day four and day twelve). Average biomass (ash free dry weight) differed between the highest and lowest biomass alga by 1.6-fold on day four and 1.7-fold on day twelve (Supplemental Figure S1). Increasing biomass between days four and twelve supports that at day eight the algae were all in mid to late exponential growth.

**Figure 1.**
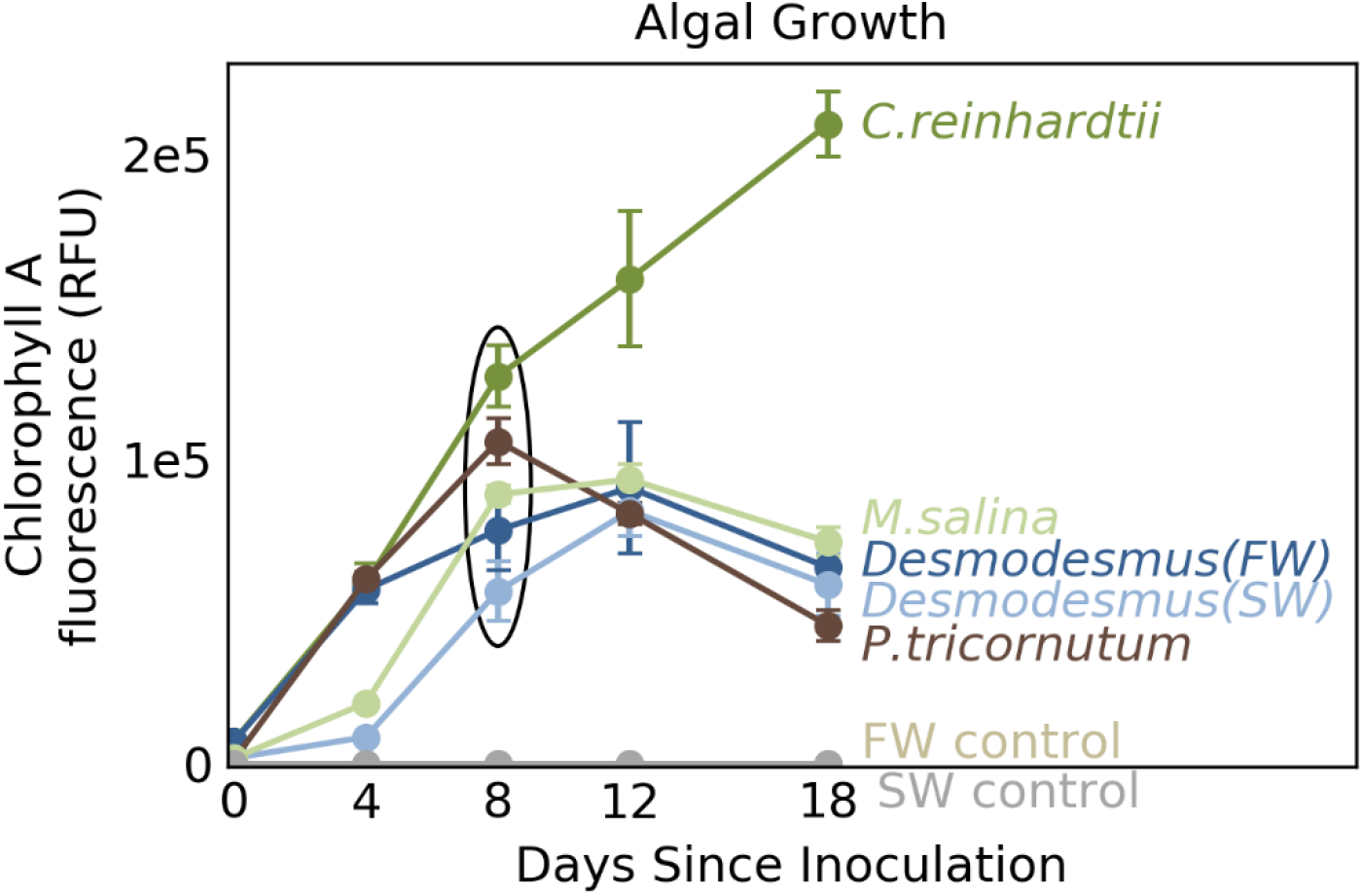
Algal growth as assessed by chlorophyll A fluorescence. Points and error bars indicate average fluorescence and standard deviation respectively of five biological replicates. The black oval indicates the day eight samples, which were used for metabolite analysis.

The total organic carbon concentration of algal spent medium was measured for samples from day eight. Organic carbon concentrations ranged from 13 ± 2 ppm (1,080 ± 170 μM) for *C. reinhardtii* to 4 ± 2 ppm (330 ± 170 μM) for *Desmodesmus* (SW) (mean ± standard deviation for five replicates) (Supplemental Figure S2). Spent medium from algae grown in freshwater (*C. reinhardtii* and *Desmodesmus* (FW)) had higher concentrations of organic carbon than spent medium from algae grown in saltwater (*P. tricornutum, M. salina*, and *Desmodesmus* (SW)). Within each growth condition, the relative concentrations of organic carbon in spent medium followed the patterns of chlorophyll A fluorescence (*C. reinhardtii* > *Desmodesmus* (FW) and *P. tricornutum* > *M. salina* > *Desmodesmus* (SW)).

### Metabolite profiles of diverse microalgae

To minimize matrix effects resulting from differences in salt concentrations, we focused on non-polar metabolite analysis using both C18-based solid phase extraction and C18-based liquid chromatography. Initial untargeted analysis of liquid chromatography tandem mass spectrometry (LC-MS/MS) data for all cell pellet, spent medium, and control samples detected 14,916 features, of which 4,259 features were determined to be significantly different from media blank controls (adjusted *p*-value < 0.05 and fold change compared to control > 2) for at least one sample group, where a sample group consisted of all five replicates for a single organism and sample type (cell pellet or spent medium) combination (Supplemental Table S1).

Within the subset of significant features, overall exometabolome composition differed between the different algal cultures (Figure 2). Principal component analyses of the exometabolite profiles separated samples from the different algae, including separating metabolite profiles for *Desmodesmus* grown in freshwater from *Desmodesmus* grown in saltwater (Figure 2). The first two principal components accounted for a total of 67.4% of the variance in exometabolite profiles. Saltwater cultures separated along the first principal component axis and freshwater cultures separated largely along the second principal component axis.

**Figure 2.**
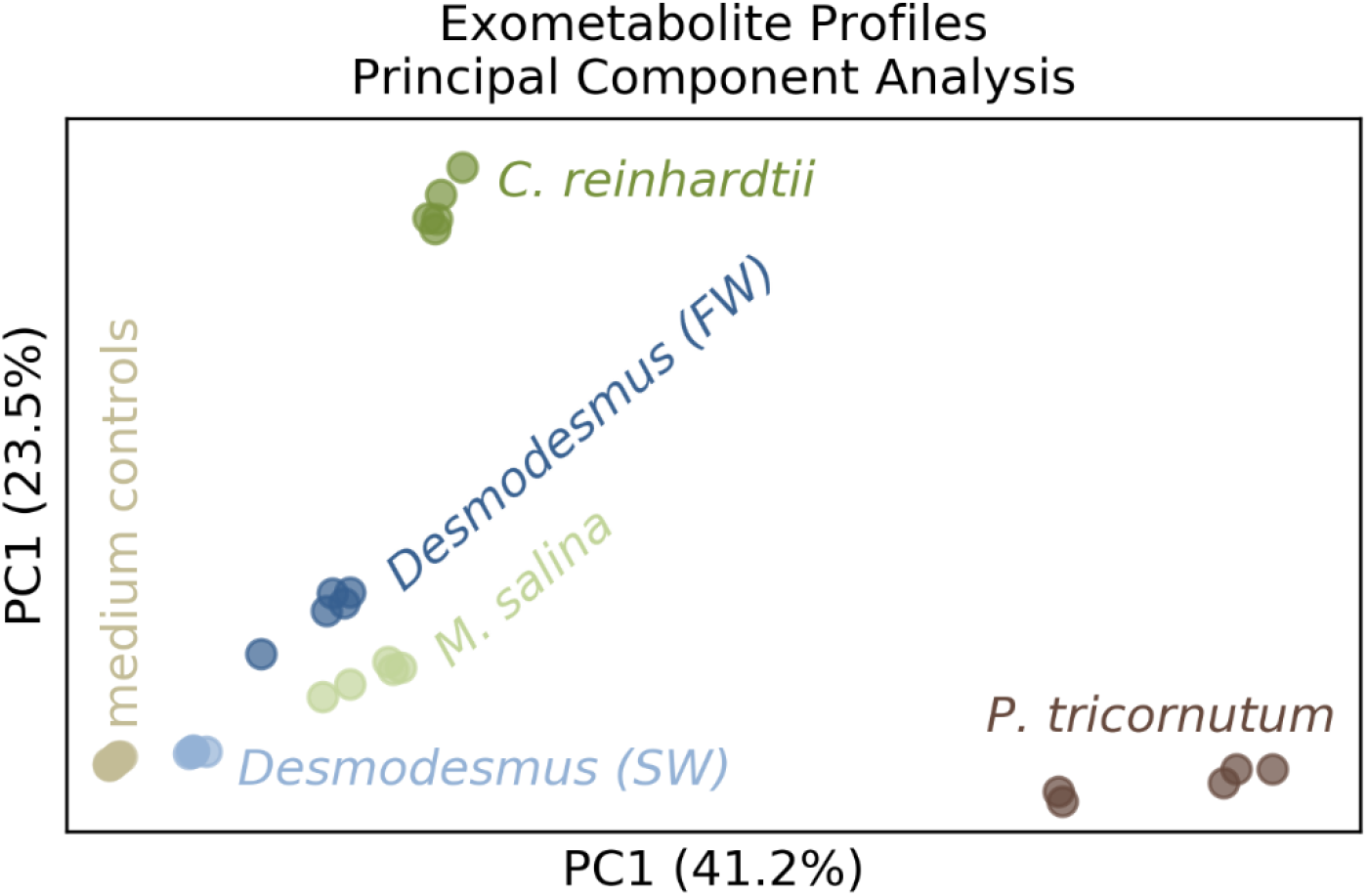
Principal component analyses of algal exometabolite profiles. Each point represents one of five replicate samples per alga or control condition. Colors indicate which algal culture the sample was from. Grey = medium controls, brown = *P. tricornutum*, light green = *M. salina*, dark green = *C. reinhardtii*, light blue = *Desmodesmus* (FW), and dark blue = *Desmodesmus* (SW).

Hierarchical clustering of LC-MS/MS (Figure 3, top dendrogram) revealed distinct sets of predominantly cell pellet features (grey) and predominantly extracellular features (purple). Hierarchical clustering of samples (Figure 3, side dendrograms) indicated that *Desmodesmus* metabolite profiles from fresh and saltwater were distinct but were more similar to each other than to metabolite profiles from other algae. Further, although *Desmodesmus* and *C. reinhardtii* are more closely related to each other than to the other algae, their metabolite profiles did not cluster together in the hierarchical clustering analysis. To further assess whether the metabolite profiles correlated with phylogeny, we conducted a Mantel test comparing phylogenetic distances between algae to Euclidean distances between normalized exometabolite profiles. That test did not detect any significant correlation between phylogeny and exometabolome (r = 0.07, *p*-value = 0.29 when the saltwater metabolite profiles were used for *Desmodesmus* and r = 0.17, *p*-value = 0.17 when the freshwater metabolite profiles were used).

**Figure 3.**
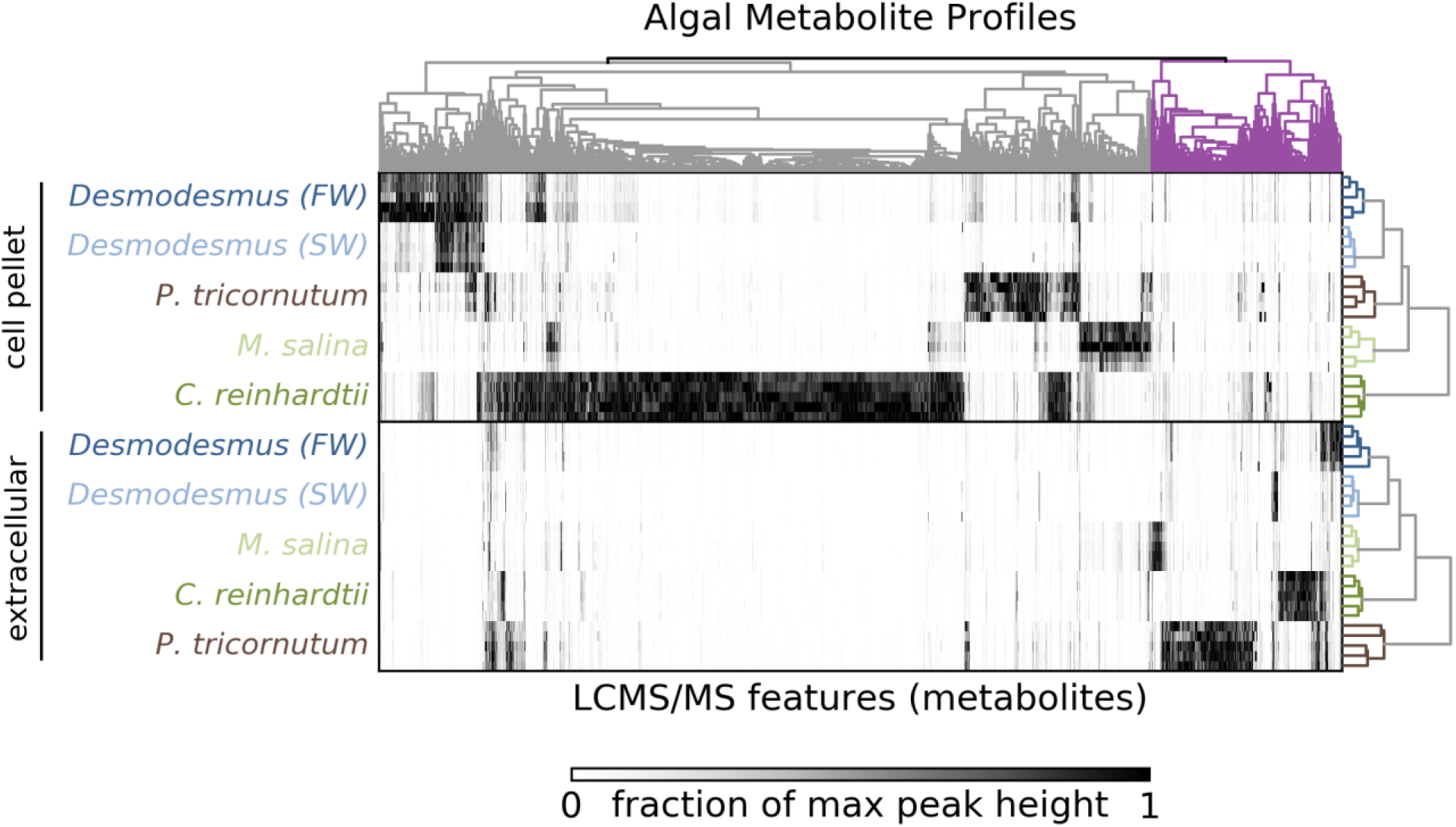
Heatmap of algal metabolite signal intensities. Samples (5 replicates per alga) are represented in rows, and metabolites are represented in columns. Shading at each point indicates the level of a feature in a particular sample as measured by the fraction of the maximum signal intensity for that feature. Top dendrogram shows hierarchical clustering of metabolites. Side dendrograms show hierarchical clustering of samples for extracellular and cell pellet samples.

To obtain putative annotations for some features and to assess potential relationships between features through feature based molecular networking, we analyzed the peak detection results with the Global Natural Products Social Molecular Networking (GNPS) tool (20). This analysis resulted in preliminary spectral matches for 263 features based on MS2 mass spectra, of which 137 (52%) had signal intensities at least twofold greater than controls for at least one sample group (Supplemental Table S2). This represents 3.2% of all 4,259 features that were significantly greater than media controls. For features of interest discussed below, we manually curated the best spectral matches (from up to 100 matches per feature) based on the GNPS cosine score, delta *m/z*, number of shared peaks, and adducts. Final curated annotations are listed in Supplemental Table S2. Only the features with the highest quality matches (GNPS cosine score > 0.7, delta *m/z* < 10 ppm, number of shared peaks > 5, common adducts: [M+H]^+^ for positive mode or [M-H]^-^ for negative mode) were considered putative annotations due to a lack of retention time information for authentic standards, with a metabolomics standards initiative (MSI) level 2 identification (21, 22). Two metabolites were subsequently confirmed using authentic standards as described below.

### Characterization of algal exudates

To differentiate metabolites resulting from lysis vs. exudation, we identified features significantly enriched in the exometabolome in paired samples of extracellular and cell pellet metabolites. Of the 138 features with spectral matches from GNPS, 41 (29%) were significantly enriched in the exometabolome of at least one alga (Figure 4).

**Figure 4.**
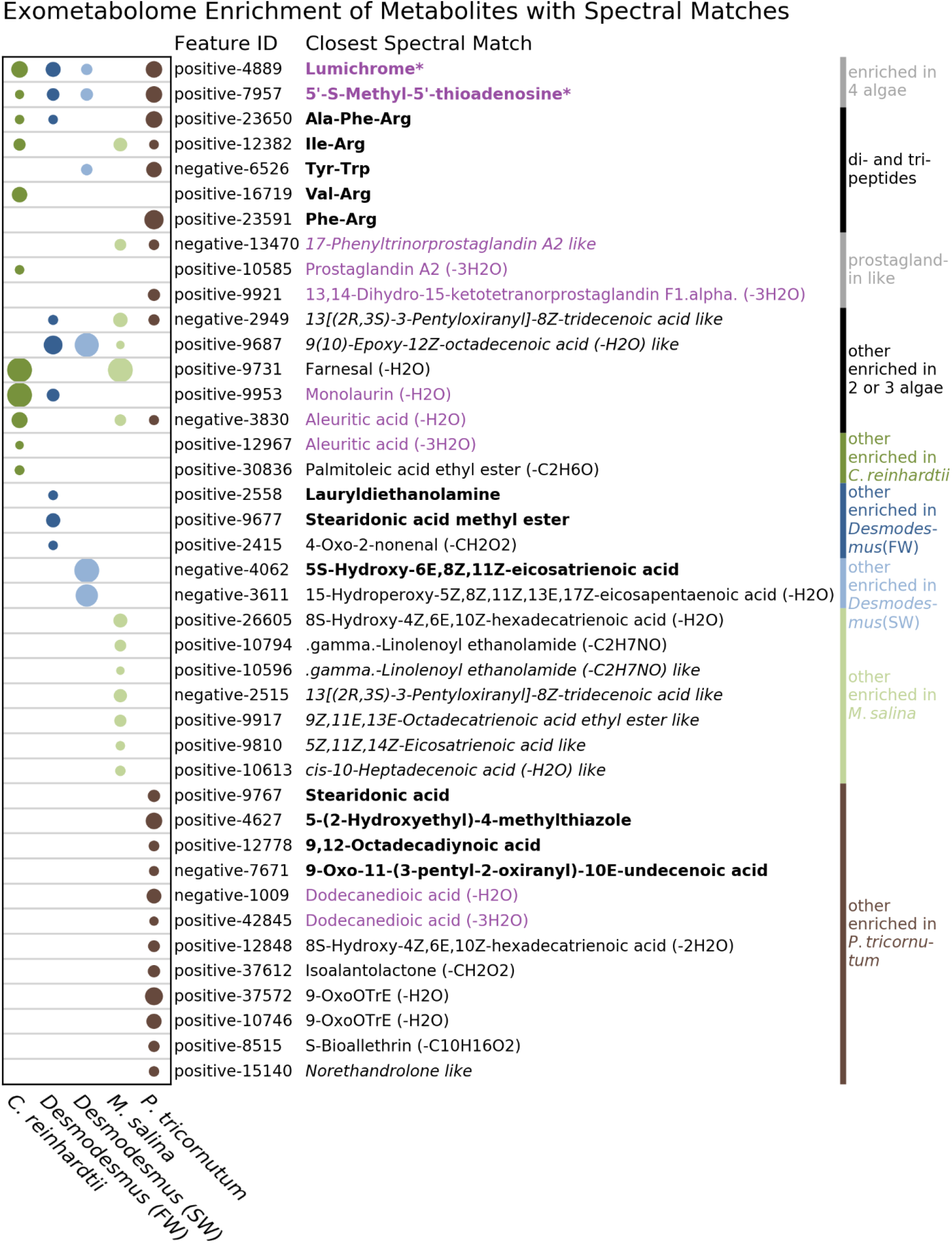
Exometabolome enriched metabolites with putative annotations. Circle sizes are proportional to the average relative enrichment in the exometabolome as compared to the cell-associated metabolome for five biological replicates. MSI level 1+ identified compounds are indicated with “*”. Spectral matches in bold indicate high quality putative annotations (MSI level 2). Spectral matches in plain text indicate quality matches to unusual adducts, suggesting related molecules with different chemical formulae, with potential difference indicated in parentheses (MSI level 3). Spectral matches in italics indicate matches with high *m/z* error, suggesting different, but potentially related compounds. Spectral match names in purple were selected for exogenous metabolite addition experiments.

Although most exometabolome enriched features were enriched in the spent medium of only one alga, two features (positive-features #4889 and #7957, identified as lumichrome and 5’-S-methyl-5’-thioadenosine, respectively) were enriched in four of the five algal systems tested. The identities of these features were further confirmed by matching retention times to standards analyzed in our laboratory, resulting in MSI level 1+ identifications. Levels of lumichrome and 5’-S-methyl-5’-thioadenosine differed between algal cultures, with *P. tricornutum* producing the highest levels of both metabolites among the exometabolomes (Supplemental Figure S3). Additionally, four features were enriched in three of the algal systems, and three were enriched in two of the algal systems (Figure 4).

Three features potentially related to prostaglandins were enriched in the exometabolomes of some algae, with at least one of these features enriched for all algal strains tested except *Desmodesmus*. Across the full dataset, seven features had best spectral matches to prostaglandins, all with either spectral matches to unusual adducts or had *m/z* errors greater than 10 ppm, indicating that these were not exact matches, but instead represented potentially related compounds. To further understand this group of features, we evaluated these features in the context of the GNPS feature based molecular networking analysis (23), which resulted in five subnetworks containing features with spectral matches to prostaglandins (Supplemental Figure S4). The features in these subnetworks were generally associated with individual algal strains, with different features being dominated by different strains. In addition to prostaglandin-like features, others had high spectral similarity to steroids and other lipids. Moreover, many of the edges in the molecular network (linking two features with similar spectra) corresponded to *m/z* differences indicating the specific loss or gain of small numbers of carbons and hydrogens (C_2_H_2_, C_4_H_4_, etc.), suggesting that this may represent a related group of lipids with small variations in chemical formulae.

Several features putatively annotated (MSI level 2) as dipeptides and tripeptides were also enriched in the exometabolomes of the different algae, with each alga having at least one of these enriched in its exometabolome. These features (positive-features #23650, #12382, #12591, and #16719, and negative-feature #6526) were putatively annotated as Ala-Phe-Arg, Ile-Arg, Phe-Arg, Val-Arg, and Tyr-Trp, respectively. Across the full dataset, di- and tripeptide features were highly associated with the exometabolomes as compared to cell pellet metabolomes, with the exception of Arg-Trp which was most abundant in the cell pellet metabolome of *C. reinhardtii* (Supplemental Figure S5).

### Impact of exudate metabolites on algal growth

Based on the enrichment of specific features in the exometabolomes of one or more algal cultures, eight metabolites were selected to evaluate their impacts on algal physiology and growth. The selected metabolites were 13,14-dihydro-15-keto-prostaglandin F1-α, 17-phenyl trinor prostaglandin A2, prostaglandin A2, 5’-S-methyl-5’-thioadenosine, lumichrome, monolaurin, aleuritic acid, and dodecanedioic acid (exometabolome enrichment of features is shown in Figure 4). Five of the added metabolites had significant and distinct effects on growth for each of the algal strains (Figure 5). Most significant was dodecanedioic acid, which impacted growth in all cultures, decreasing growth of *C. reinhardtii, Desmodesmus* (FW) and *P. tricornutum* (65%, 14%, and 23% respectively), and increasing growth of *Desmodesmus* (SW) and *M. salina* (44% and 48% respectively, Figure 5). Aleuritic acid had opposite effects on the two freshwater cultures, increasing *C. reinhardtii* growth and decreasing *Desmodesmus* freshwater growth (Figure 5). Lumichrome, 5’-S-methyl-5’-thioadenosine, and 17-phenyl trinor prostaglandin A2 increased growth in *P. tricornutum* (17%, 19%, and 20% respectively) and the latter two decreased growth in *Desmodesmus* in freshwater.

**Figure 5.**
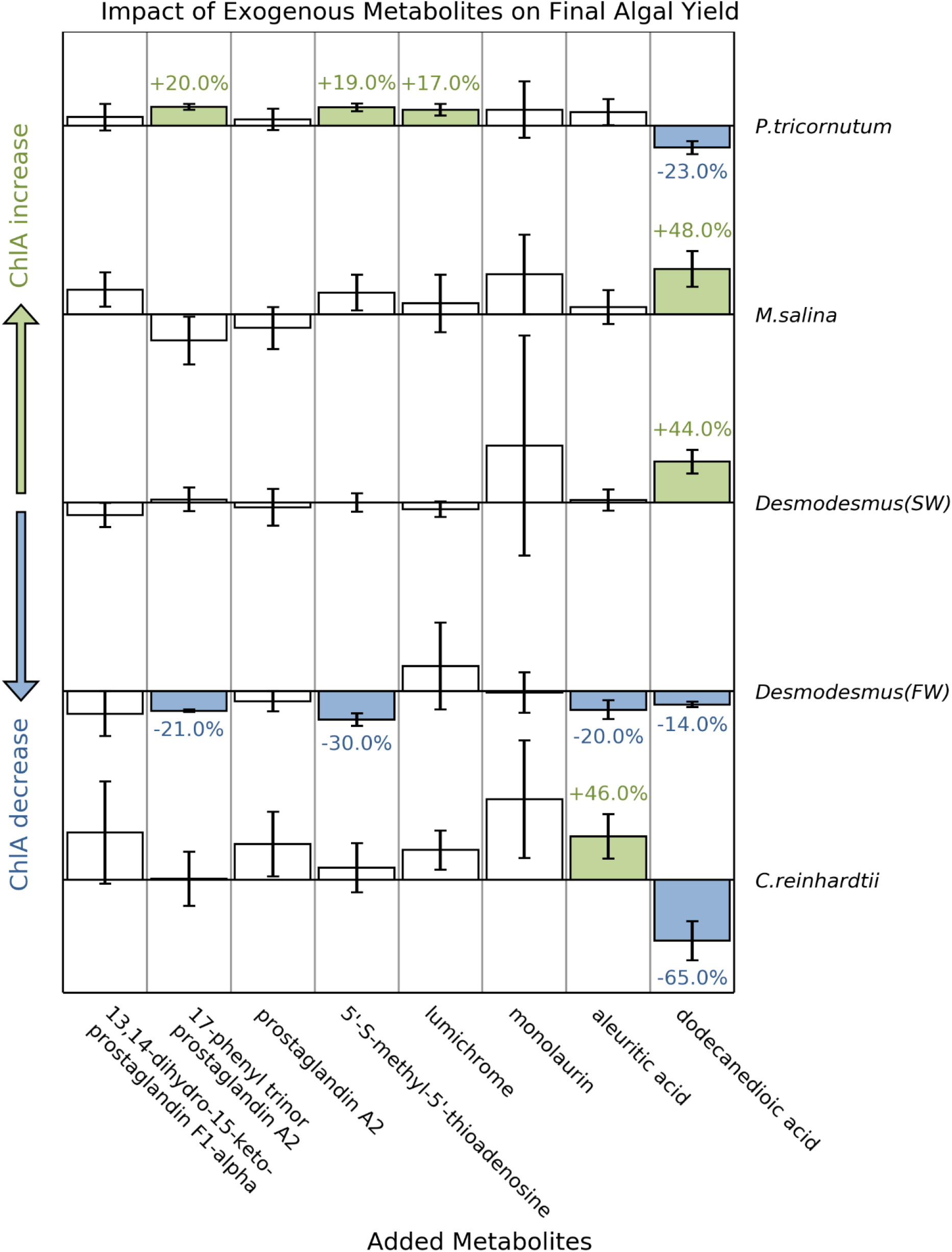
Impacts of select metabolites on algal growth as measured by chlorophyll A fluorescence. Bars and error bars show the average and standard deviation of the increase or decrease of chlorophyll A fluorescence relative to controls of three biological replicates. Filled bars indicate a statistically significant decrease (blue) or increase (green), with the percent increase or decrease indicated next to bars.

### Response of *Desmodesmus* to media conditions

*Desmodesmus* sp. strain C406 is unique among the algae tested here in its ability to grow in both the saltwater and freshwater medium. Between the two growth media, both the extracellular and cell pellet metabolite profiles differed significantly. Because high salt concentrations can interfere with detection of compounds by LC-MS/MS, making some metabolites appear to be lower or not detectable from high salt samples due to matrix effects (24, 25), we focused on metabolites that were detected at significantly higher levels in the saltwater samples. Three features with spectral matches to database compounds were significantly enriched in the exometabolome of *Desmodesmus* grown in saltwater compared freshwater (Supplemental Figure S6). These enriched features included the putatively annotated dipeptides Tyr-Trp and Arg-Trp, as well as negative-feature #1013, which had high spectral similarity to the [M-H-H_2_O]^-^ adduct of dodecanedioic acid. Several of the metabolites resulted in decreased growth in *Desmodesmus* (FW), but did not have significant effects on *Desmodesmus* (SW). Notably, in the exogenous metabolite addition experiment above, *Desmodesmus* responded positively (44% increase in chlorophyll A) to dodecanedioic acid when grown in saltwater, but negatively when grown in freshwater (14% decrease in chlorophyll A) (Figure 5).

## DISCUSSION

Our study builds on previous work to expand our understanding of microalgal exometabolome diversity, composition, and impacts on algal interactions. We set out to characterize exometabolomes from a phylogenetically and ecologically diverse set of microalgae including two green algae (one freshwater, one freshwater and marine), a marine diatom and a marine eustigmatophycean alga to discover exometabolites that may mediate algal interactions and ecophysiology. Given our interest in comparing algal responses to different salinities, we focused our analysis on non-polar metabolites and used solid phase extraction to minimize the influence of salinity on metabolite analysis. We find that (A) exometabolite profiles are distinct between these eukaryotic microalgal strains but we did not detect any correlation of exometabolomes with phylogeny, (B) based on the distinct responses to exogenous addition and salt and freshwater culturing, exometabolite profiles may instead primarily reflect algal ecophysiology, (C) lumichrome, dipeptides, and prostaglandin-like compounds were enriched in the algal exometabolomes, none of which are well characterized in algal systems, and (D) several compounds acted as effector metabolites, having significant impacts on algal growth. The exometabolites profiled may play roles in growth promotion, defense, and interactions as has been shown in other systems.

In our comparison across phylogenetically diverse algal strains, we found distinct metabolite profiles associated with each strain. One previous study has compared the exometabolomes of two cyanobacterial groups (three *Prochlorococcus* strains and two *Synechococcus* strain) with diatoms (two *Thalassosira* species and one *Phaeodactylum*) (16). That study found that the differences in the exometabolome profiles reflected the phylogenetic differences (prokaryote vs eukaryote) between these groups. Our results indicate that this phylogenetic signal is not consistently present within eukaryotic microalgae. *C. reinhardtii* and *Desmodesmu*s, both members of the phylum Chlorophyta and class Chlorophyceae, are more closely related than the other strains used in our study (a diatom and a eustigmatophycean alga). However, their metabolite profiles did not cluster together in our analysis (Figure 3, side dendrograms). Further, Mantel tests did not detect significant correlation between algal phylogenetic distance and exometabolome composition distance. A study of both macro and microalgae found that environmental origin influenced intracellular metabolome composition more than phylogenetic relatedness (26). Our study suggests that exometabolite profiles are also not phylogenetically determined.

The distinct profiles and response of *Desmodesmus* under fresh and saltwater conditions as well as the distinct responses of the microalgae to the exogenously added metabolites suggests that algal ecophysiology may be an important determinant of exudation. Given the potential signal suppression at high vs. low salt concentrations (even using solid phase extraction) we focus on metabolites that increased with high salt. We find that *Desmodesmus* drastically shifts its exometabolome in response to salt and freshwater culturing and has contrasting growth responses to addition of three of the eight tested metabolites under the different conditions, highlighting the responsiveness of the exometabolome to shifting environmental conditions. There have been some studies examining intracellular responses to different salinities by a variety of microalga including *Scenedesmus* sp., *Amphora subtropica, Dunaliella* sp., *Desmodesmus armatus, Mesotaenium* sp., and *Tetraedron* sp (27–30). Our work supports the finding that salinity impacts algal metabolite composition and extends it to the extracellular environment, suggesting that these differences will influence the surrounding community as well.

Several of the exometabolome enriched metabolites may act as effector metabolites, with potentially important roles in algal growth and interactions with bacteria. Lumichrome (MSI identification level 1+), which was enriched in the exometabolomes of all cultures except *M. salina*, has been found to promote plant development and growth and has also been shown to impact bacterial quorum sensing (31, 32). *Chlamydomonas* is known to produce lumichrome (31), but lumichrome production has not been reported previously in *P. tricornutum* or *Desmodesmus*. Most previous studies of algae and lumichrome or riboflavin (vitamin B2), of which lumichrome is a derivative, have focused on possible lumichrome/riboflavin production by algal growth promoting bacteria, as opposed to algal production (33–35). Two recent studies found that exogenous lumichrome had positive growth impacts on the green algae *Chlorella sorokiniana* and *Auxenochlorella protothecoides* (33, 34), while another found that riboflavin increased chlorophyll production in *Chlorella vulgaris* (35). Supporting these studies, we found that *P. tricornutum*, which produced the highest level of lumichrome (Supplemental Figure S3), exhibited increased growth in response to exogenous lumichrome addition (Figure 5), indicating it may be particularly important for this model diatom. To our knowledge this is the first study showing impacts of lumichrome on a diatom or any saltwater microalga, expanding the range of systems where this compound may have important impacts. Further, given that lumichrome was enriched in four of the five exometabolomes and its role in bacterial quorum sensing, it could also play a role in modulating the phycosphere bacterial community.

Prostaglandin like compounds were another group which were enriched in the exometabolomes of several of the algal strains (*C. reinhardtii, P. tricornutum*, and *M. salina*). Our data suggests that these features may represent related but novel compounds because these spectral matches were to unusual adducts or had a *m/z* error greater than 10 ppm. Prostaglandins are hormone like signaling molecules found in a range of animals and plants (36–38). Originally discovered and studied in animals, prostaglandins have only recently been detected in microalgae, first being reported in the diatom *Skeletonema marinoi* (37). Although the roles of these compounds in microalgae have not yet been elucidated, these molecules are known to be involved in intracellular communication in other organisms. Prostaglandins have also been found to have diverse roles in marine macroalgae, including as defense molecules against pathogens in red macroalgae and as part of the oxidative response in brown macroalgae (36). A recent study in *P. tricornutum* detected the production of the isoprostanoids, which are structurally related to prostaglandins, under oxidative stress (39). That study found that the addition of 5 to 50 μM of several different isoprostanoids to *P. tricornutum* cultures decreased cell counts and increased lipid accumulation, while not affecting photosynthetic efficiency. In comparison, we found that the addition of 10 μM of 17-phenyl trinor prostaglandin A2 increased chlorophyll A production by *P. tricornutum* by 20%, and decreased chlorophyll production of *Desmodesmus* grown in freshwater but not saltwater (Figure 5). Interestingly, negative-feature #13470, which showed spectral similarity to 17-phenyl trinor prostaglandin A2, was enriched in the exometabolome of *P. tricornutum*, but not that of *Desmodesmus* (Figure 4). Together, these results suggest that prostaglandins and related compounds have important roles affecting microalgal growth.

Another interesting group of detected compounds included the di- and tripeptides, of which at least one was enriched in each of the five algal system exometabolomes profiled, indicating that this group of exometabolites is widespread across diverse microalgae. We identified one tripeptide and several dipeptides enriched in the alga exometabolomes, with specific dipeptides highly associated with either *P. tricornutum* or *C. reinhardtii* (Supplemental Figure S5). Dipeptides are known to be involved in inter-kingdom interactions as well as bacterial quorum sensing. For example, dipeptides have been implicated in coral-algal symbiosis (40, 41), as well as the interactions between land plants and arbuscular mychorhizal fungi (42). Cyclic dipeptides are important compounds for quorum sensing in Gram-positive bacteria and are also produced by other organisms to interfere with that quorum sensing (43). Linear dipeptides, like those detected in our study, have also been found to affect bacterial quorum sensing systems (44). The specificity of these dipeptides to particular algal taxa suggests that these metabolites may not be the result of general proteolysis, but instead that these particular peptides may have important roles for these algae. Because these metabolites are enriched in the exometabolome, they could play a role in inter-kingdom interactions, as has been seen in these other systems.

The contrasting growth responses to exogenous addition of specific effector metabolites suggest there is delicate and distinct homeostasis with respect to these compounds and their extracellular and intracellular concentrations. Some metabolites are excreted to reduce toxicity but can also have growth promoting effects. Indole-3-acetic acid, a plant hormone also produced by some algae, stimulates growth of *Desmodesmus* at low concentrations, but inhibits growth at concentrations above 200 μM (45). Contrasting growth responses of different algae to the same metabolite could also indicate alleopathy, in which one alga releases metabolites to inhibit competitors. In our study, negative-feature #3830, with spectral similarity to the [M-H-H_2_O]^-^ adduct of aleuritic acid, was enriched in the exometabolomes of *C. reinhardtii*, *M. salina*, and *P. tricornutum*, but not in the exometabolome of *Desmodesmus* (Figure 4). In the growth impacts experiment, aleuritic acid inhibited the growth of *Desmodesmus* but not the other algae (Figure 5).

There are some limitations to this study which are important to consider. First, our analysis has focused on non-polar metabolites, which enabled analysis of changes in response to salt concentration and analysis of compounds relevant to signaling. However, excluding polar metabolites provides a limited view of overall exudates. Second, given this focus on diverse non-polar metabolites relatively few were identified. Therefore, an important future direction will be the extension of this work to the analysis of polar metabolites which include many well-characterized molecules such as sugars, amino acids, and small organic acids. Further, given the large phylogenetic distances and limited number of taxa profiled, there are some limitations to what can be concluded about phylogenetic correlations and the specificity of exometabolites to phyla. However, because of the wide diversity of the algal systems studied, the similarities provide evidence for exometabolites that are commonly exuded by microalgale

This work adds to our understanding of microalgal exometabolomes. We found that variation in eukaryotic microalgal exometabolome composition is not primarily phylogenetically driven and is affected by growth conditions. We also identified impacts of specific effector metabolites, many which have not been identified or characterized in algae, on algal growth. Further work is now needed to understand the roles of these metabolites in more complex algal-bacterial systems and to extend this work to include polar metabolites.

## MATERIALS AND METHODS

### Algal strains and growth conditions

This study used four microalgal strains, as detailed in Table 1. Two of the algal strains (*C. reinhardtii* and *Desmodesmus*) were grown in freshwater medium, and three (*P. tricornutum*, *M. salina*, and *Desmodesmus*) were grown in saltwater medium.

**Table 1.** Growth media, organisms, and strain information for algal growth experiments.

| Medium | Organism | Strain |
| --- | --- | --- |
| Freshwater<br>(Bristol’s medium) | *Chlamydomonas reinhardtii* | CC-1690 |
|  | *Desmodesmus* sp. | C046 |
| Saltwater<br>(ESAW medium) | *Phaeodactylum tricornutum* | CCMP 2561 |
|  | *Microchloropsis salina* | CCMP 1776 |
|  | *Desmodesmus* sp. | C046 |

Saltwater growth medium was prepared using salts for Enriched Saltwater Artificial Water (ESAW) medium (Berges et at 2001, Harrison et al. 1980). Saltwater medium contained 21.194 g/L NaCl, 3.55 g/L M=Na_2_SO_4_, 0.599 g/L KCl, 0.174 g/L NaHCO_3_, 0.0863 g/l KBr, 0.023 g/L H_3_PO_3_, 0.0028 g/L NaF, 9.592 g/L H=MgCl_2_·6H_2_O, 1.344 g/L CaCl_2_·2H_2_O, 0.0218 g/L SrCl_2_·6H_2_O, and 0.03 g/L Na_2_SiO_3_·9H_2_O. Additionally, nitrate, phosphate, trace metals, and vitamins were added at the concentrations for F/2 medium (46). Nitrate and phosphate were added as 0.075 g/L NaNO_3_ and 0.005 g/L NaH_2_PO_4_.H_2_O respectively. Trace metals final concentrations were 0.023 mg/L ZnSO_4_·7H_2_O, 0.152 mg/L MnSO_4_·H_2_O, 0.0073 mg/L NaMoO_4_·2H_2_O, 0.014 mg/L CoSO_4_·7H_2_O, 0.0068 mg/L CuCl_2_·2H_2_O, 4.6 mg/L Fe(NH_4_)2(SO_4_)_2_·6H_2_O, and 4.4 mg/L Na_2_EDTA·2H_2_O. Vitamin final concentrations were 0.0135 mg/L cyanocobalamin (Vitamin B12), 0.0025 mg/L biotin, and 0.0335 mg/L thiamine.

Freshwater medium was prepared based on Bristol’s medium. The medium contained 0.025 g/L NaCl, 0.025 g/L CaCl_2_·2H2O, 0.075 g/L MgSO_4_·7H_2_O, 0.25g NaNO_3_, 0.075g/L K_2_HPO_4_, and 0.175 g/L KH_2_PO_4_. To make the freshwater and saltwater conditions more comparable, the same F/2 levels of trace metals and vitamins listed above for saltwater medium were also added to the freshwater medium.

Algal were grown in glass 125 mL Erlenmeyer flasks that had previously been baked in a muffle furnace at 550 °C for two hours to remove trace organics that could contaminate metabolomics analyses. Each flask contained 50 mL of the appropriate medium and was capped with a foam stopper and covered with aluminum foil to avoid contamination during the experiment. Flasks were incubated with shaking at 90 rpm in a light incubator, with a 12-hour day/night light cycle. Daytime illumination was 3500 lux, and the incubator temperature was maintained at 22 °C. Algal inocula were prepared by first growing algal stock cultures under experimental conditions (flasks, media, incubation) for one week to acclimate cultures to those conditions. After seven days, experimental flasks were inoculated by transferring 2 mL of inoculum culture to flasks with the appropriate medium. Twenty-five biological replicate cultures were prepared for each algal and control condition to allow destructive sampling of five replicates at each sampling time point (0, 4, 8, 12, and 18 days).

### Sample collection

Chlorophyll A measurement samples were collected by transferring 1 mL of culture to a clear 5 mL polystyrene tube. Microscopy samples were collected by transferring a 1 mL sample from the flask to a 1.5 mL microcentrifuge tube, adding 100 μL of formaldehyde to fix cells, and storing the sample at 4 °C until performing microscopy to determine axenicity of the culture.

After removal of samples for chlorophyll measurements and microscopy, the remainder of the culture was destructively harvested to collect cell pellets and spent medium. Spent medium and cell pellets were separated by centrifugation. Half of the culture was transferred to a 50 mL falcon tube and centrifuged at 5000 g for eight minutes at 4 °C. The supernatant was carefully transferred to a clean Falcon tube, and the remainder of the culture was added to the tube containing the pellet and the centrifugation procedure was repeated. After separation by centrifugation, the supernatant was filtered through a 0.45μm pore size syringe filter to remove residual algal cells. Filtered spent medium and cell pellets were frozen on dry ice and stored at −80 °C until extraction.

In order to disrupt cells for metabolite extraction, cell pellets were resuspended in 1 mL sterile water, frozen at −80 °C, and lyophilized. Lyophilized pellets were disrupted using a sterilized steel ball and vortexing three times for five seconds. Disrupted pellets were then resuspended in 50 mL sterile water and filtered and frozen in the same manner as spent medium. Thus, the spent medium and cell pellet samples corresponded to the same original sample volume.

### Chlorophyll A Fluorescence, Biomass, and Organic Carbon Measurements

Chlorophyll A fluorescence of samples from flask grow cultures was measured using a Trilogy Fluorometer [Turner Designes, San Jose, CA, USA] with the Chlorophyll A In-Vivo Module.

Biomass was measured as the ash free dry weight of the cell pellets collected on days four and eight. Cell pellets were transferred to aluminum dishes and dried at 95 °C over night. The dry weight of the pellet and aluminum dish was measured. Pellets were then ashed at 500 °C for four hours, and the weight of the remaining ash and aluminum dish was measured. Ash free dry weight was calculated as the difference between the two measured weights.

Samples were prepared for total organic carbon (TOC) measurement by diluting 1.4 mL sample (spent medium or cell pellet extract) with 5.6 mL ultrapure water for a 1:5 dilution. TOC was measured on a Shimadzu TOC analyzer.

### Metabolite profiling

Samples for LC-MS/MS metabolomics analysis were extracted by solid phase extraction using Waters Sep-Pak C18 3 mL cartridges. Because chlorophyll and biomass content were similar across samples, samples were normalized to the same volume (30 mL). Cartridges were first preconditioned with 3 mL methanol, and the rinsed with 6 mL ultrapure water. Metabolites were extracted by passing 30 mL of sample through the cartridge. Columns with extracted metabolites were rinsed with 6 mL ultrapure water. Metabolites were eluted with 1 mL methanol into a 1.5 mL polypropylene tube and the dried in a vacuum centrifuge with a refrigerated vapor trap. Dried metabolite samples were resuspended in 150 μL methanol with matrix control internal standards (Supplemental Table S3), vortexed for 5 to 10 seconds, and sonicated in a water bath sonicator with ice for ten minutes. Samples were centrifuged at 10,000g for five minutes at 4 °C to pellet insoluble salts and proteins. Supernatant was transferred to a 0.2 μm pore size centrifuge filter device, and centrifuged at 5,000 g for five minutes at 4 °C. Filtrate was transferred to an autosampler vial for LC-MS/MS analysis.

LC-MS/MS was performed as previously described (47), and the LC gradient and MS conditions are provided in Supplemental Table S4. Briefly, chromatographic separation of metabolites in samples were performed using reverse-phase liquid chromatography on a ZORBAX EclipsePlusC18 RRHD 1.8 μm, 2.1 x 50 mm column [Agilent Technologies, Santa Clara, CA, USA] on an Agilent 1290 series ultra high-performance LC system. Mass spectrometry analyses were performed using a Thermo Fisher Scientific Q Exactive Hybrid Quadrupole-Orbitrap Mass Spectrometer [Thermo Fisher Scientific, Waltham, MA, USA]. LCMS parameters used for analysis are defined in Supplemental Table S4.

Untargeted analysis was LC-MS/MS data was performed to identify metabolites as previously described (47). Briefly, LC-MS/MS features, corresponding to specific retention time and m/z combinations, were detected using the MZMine software v. 2.53 (48, 49). Detected features were analyzed using the Global Natural Products Social Molecular Networking tool (GNPS) to construct feature based molecular networks (23, 50). This analysis creates networks of potentially related features, using spectral similarity as a proxy for structural similarity. Preliminary feature annotations were determined based on the highest scoring MS2 match to the GNPS database. For features identified as statistically significant in subsequent analyses, putative annotations were made by hand curation of the top 100 matches to the GNPS database, based on GNPS cosine score, delta *m/z*, number of shared peaks, and common adducts.

### Impacts of select metabolites on algae

Impacts of selected metabolites on algal chlorophyll A production were tested in 48 well plates. Each metabolite was tested individually for each of the algal strains at a concentration of 0.01 mM. Each well contained 750 μL of media with or without added metabolite and was inoculated with 15 μL of one week old algal inoculum. Each metabolite was tested for each alga in triplicate in randomly positioned wells. Plates were incubated with a 12-hour light/dark cycle for 14 days with shaking at 90 rpm. Chlorophyll A fluorescence of plate growth assays was measured in a Cytation5 plate reader [BioTek, Winooski, VT, USA] with excitation and emission wavelengths of 440 nm and 680 nm respectively.

### Statistical analyses

LC/MS-MS features significantly different from background were identified by comparing feature peak heights for test samples to those of media blanks with two tailed t-tests and using the Bonferroni correction to account for multiple comparisons. Features were considered significant if at least one sample group had an adjusted *p*-value < 0.05 and at least 2-fold above the signal in control samples.

Features significantly enriched in the exometabolome were identified by first calculating the ratio of the peak heights for paired exometabolome and pellet metabolome samples. A one sample t-test was used to compare mean of the base two logarithm of these ratios to zero, using the Benjamini-Hochberg adjustment to correct for multiple comparisons. A feature was considered significantly exometabolome enriched for a particular alga if the adjusted *p*-value was less than 0.05 and average ratio of extracellular to cell pellet peak height was at least 2.

Mantel tests were conducted using the “mantel” function in the “vegan” package v. 2.5-7 in R v. 4.0.5. Phylogenetic distances were calculated based on multiple sequence alignment of 18S rRNA genes to the global SILVA alignment for rRNA genes (51). Metabolome distance matrices were determined based on Euclidean distances of normalized metabolite profiles. Separate Mantel tests were performed using either the freshwater or saltwater exometabolomes for *Desmodesmus* in the metabolome distance matrix.

## Data availability

GNPS molecular networks and preliminary spectral matches are available at https://gnps.ucsd.edu/ProteoSAFe/status.jsp?task=29549d19521944368add9234d215fa1b and https://gnps.ucsd.edu/ProteoSAFe/status.jsp?task=f14b47591e6c40aab59907aa2f631059. The GNPS expanded spectral matches used for curation of feature annotations are available at https://gnps.ucsd.edu/ProteoSAFe/status.jsp?task=42f8bb1af9f74575b985da04fd1e6c77 and https://gnps.ucsd.edu/ProteoSAFe/status.jsp?task=19fd0e195daa481893aa7e6f93de4573.

## ACKNOWLEDGEMENTS

This research was supported by the LLNL Biofuels Scientific Focus Area, funded by the U.S. Department of Energy Office of Science, Office of Biological and Environmental Research Genomic Science program under FWP SCW1039. This work was performed under the auspices of the U.S. Department of Energy by Lawrence Livermore National Laboratory under Contract DE-AC52-07NA27344. LLNL IM release number LLNL-JRNL-823908-DRAFT.

## SUPPLEMENTAL FIGURE CAPTIONS

**Supplemental Figure S1.** Algal biomass as measured by ash free dry weight for a 50 mL culture volume. Points indicate average value, and error bars represent standard deviation of five biological replicates.

**Supplemental Figure S2.** Total organic carbon of spent medium from eight-day old algal cultures. Bar heights indicate average value, and error bars indicate standard deviation of five biological replicates.

**Supplemental Figure S3.** Abundance of lumichrome and 5’-S’methyl-5’-thioadenosine in algal exometabolomes. These feature identifications, initially based on high quality spectral matches, were further confirmed by matching retention times to laboratory prepared standards (MSI level 1 identifications).

**Supplemental Figure S4.** GNPS molecular network subgraphs containing features with spectral matches to prostaglandins. Network edges indicate spectral similarity between LC-MS/MS features (nodes). Chemical formulae on edges indicate the difference in chemical formulae between connected features based on *m/z* differences. Pie charts indicate average relative abundance for five biological replicates of the feature among the different algal strains. Pie charts are only plotted for features that are at least twofold above background for at least one sample group. Features with spectral matches are labeled with feature IDs and, and best spectral matches are listed in the box on the lower right. Spectral matches in bold indicate high quality putative annotations (MSI level 2). Spectral matches in plain text indicate quality matches to unusual adducts, suggesting related molecules with different chemical formulae, with potential difference indicated in parentheses (MSI level 3). Spectral matches in italics indicate matches with high *m/z* error, suggesting different, but potentially related compounds (MSI level 3).

**Supplemental Figure S5.** Heatmap of putatively identified di- and tripeptides across sample types and organisms. All features are putative annotations (MSI level 2) based on high quality spectral matches of LC-MS/MS features to database reference spectra. Samples (5 replicates per alga) are represented in rows, and individual di- and tripeptide features are represented in columns. Shading at each point indicates the level of a feature in a particular sample as measured by the fraction of the maximum signal intensity for that feature.

**Supplemental Figure S6.** Metabolites enriched in the exometabolome of *Desmodesmus* under saltwater conditions as compared to freshwater conditions. Bold lettering indicates that these are putative annotations based on high quality spectral matches of LC-MS/MS features to database reference spectra (MSI level 2 annotations). Unformatted lettering indicates that this feature had a high-quality spectral match to an unusual adduct, with the unusual part of the adduct indicated in parentheses (MSI level 3 annotation).

## SUPPLEMENTAL TABLE CAPTIONS

**Supplemental Table S1.** LCMS feature peak heights. Features that were significantly different from controls are presented first and are highlighted in green.

**Supplemental Table S2.** LC/MS-MS features with spectral matches in GNPS. Curates best spectral matches were determined only for features that were identified as statistically significant in one of the analyses discussed in the paper. Curated best spectral matches in bold indicate high quality putative identifications based on high MQ score, low *m/z* error, high number of shared peaks, and common adducts. Curated best spectral matches in unformatted text indicate quality matches to unusual adducts, suggesting related molecules with different chemical formulae, with potential difference indicated in parentheses. Curated best spectral matches in italics indicate matches with high *m/z* error, suggesting different, but potentially related compounds. An “X” indicates features that were detected in the pellet or spent medium for each alga.

**Supplemental Table S3.** LC-MS/MS internal standards

**Supplemental Table S4.** LCMS instrument and parameters for metabolomics analysis.

